# Co-Substrate Free Valorisation of Lignin Monomers by Assimilation of C1 and C2 By-Products

**DOI:** 10.1101/2025.02.17.638771

**Authors:** Daniel Bergen, Òscar Puiggené, Esteban Marcellin, Robert E. Speight, Pablo I. Nikel, Birgitta E. Ebert

## Abstract

Lignin is an underutilised global resource with significant potential for the production of chemicals that are currently derived from fossil resources. However, biotechnological lignin valorisation faces various challenges including its recalcitrance and the toxicity of aromatic intermediates, products, and by-products like formaldehyde. While biochemical production from lignin-derived monomers has been demonstrated by disrupting native lignin degradation pathways, this approach required co-feeding additional carbon sources such as glucose for growth. This dependence on additional carbon sources can create competition with the food industry and undermine the economic sustainability of the bioprocess. Here, we report growth of a protocatechuate production strain of *Pseudomonas putida* EM42 on the by-products from *p*-coumarate and ferulate valorisation, achieving carbon efficiencies of up to 78 %. Additional flux balance analysis identified C1 assimilation pathways, including two novel pathways, beneficial for the growth on the formaldehyde by-product from ferulate degradation leading to improved carbon utilisation. This study demonstrates how by-product utilisation from lignin conversion can eliminate the need for co-feeding additional carbon sources, thereby potentially improving the efficiency of lignin valorisation.

## Introduction

Comprising 15-40 % of dry weight in terrestrial plants, lignin is the second most abundant biopolymer worldwide (Ragauskas *et al*., 2014). Despite its abundance, the heterogenous nature of lignin limits its use as a resource for chemical production. Chemically depolymerised lignin is a highly complex mixture of various aromatic compounds (Becker & Wittmann, 2019; Rinaldi *et al*., 2016), from which individual compounds are difficult to fractionate. Biotechnological processes are better suited to convert such heterogenous mixtures (Kamimura *et al*., 2017; Linger *et al*., 2014; Weiland *et al*., 2022). This process captures a range of lignin monomers and converts them into a single compound. This concept, called metabolic funnelling, allows for the targeted production of value-added products from complex feedstocks.

Despite progress, current biological lignin valorisation processes face challenges, in particular its reliance on co-feeding other carbon sources for biomass formation, e.g. glucose or acetate (Becker *et al*., 2018; Johnson *et al*., 2017; Kohlstedt *et al*., 2018; Rouches *et al*., 2021; Werner *et al*., 2023). This dependency is particularly pronounced in strains engineered to accumulate valuable intermediates from aromatic metabolism, as these strains are unable to fully utilise aromatic substrates as carbon and energy sources. Supplementing additional carbon sources is effective in maintaining growth of these engineered microbial strains. However, apart from increasing production costs, incorporating edible carbon sources undermines the advantage of using lignin, which is inedible and does not compete with food supply chains. Competition with the food industry is especially of growing concern given the increasing global population.

Efforts to achieve “glucose-free” lignin valorisation, where lignin monomer fractions (H-, G-, S- lignin monomers) are bifurcated into substrates for growth and precursors for production (Almqvist *et al*., 2021; Shinoda *et al*., 2019), avoid co-substrate supplementation but minimise the carbon conservation of lignin monomers in target products. Previous studies discuss growth on the C2 by-product fraction from *p*-coumarate (Fenster et al., 2022; Salvachúa *et al*., 2018). However, as far as we are aware, no study has investigated whether the acetyl-CoA (C2) by-product fraction from lignin monomers, such as *p-*coumarate and ferulate, is sufficient as sole carbon source to sustain growth of lignin valorising production strains.

C1 molecules, such as formaldehyde, also occur as by-products during aromatic degradation, e.g. methoxylated G- and S-lignin monomers (Klein *et al*., 2022). Considering that the C2 by-product fraction might be insufficient as sole carbon source for proper growth of the microbial host, as suggested by Salvachúa *et al*., (2018), enabling C1 assimilation might be a promising addition to support growth on G- and S-monomers (e.g., ferulate, vanillin, vanillate, syringate). Such endeavours could harness synthetic pathways primarily developed for the utilisation of CO_2_-derived molecules as carbon feed for biotechnological processes (Bruinsma *et al*., 2023; He *et al*., 2020; Turlin *et al*., 2022; Wenk *et al*., 2022).

This study explores the potential of using the C1 and C2 by-product fractions from the degradation pathways of *p*-coumarate and ferulate for growth and cell maintenance while conserving the core aromatic structure in the target product. *Pseudomonas putida* was chosen as production host due to its native lignin degradation pathways (Harwood & Parales, 1996; Jiménez *et al*., 2002; Nogales *et al*., 2017). Furthermore, *Pseudomonas* possess high tolerance towards oxidative stress and cytotoxic compounds as exhibited by aromatic compounds such as lignin monomers (Blank *et al*., 2008; Ebert *et al*., 2011; Nikel *et al*., 2016, 2021), while being genetically accessible with a variety of genetic engineering tools (Neves *et al*., 2020, 2024; Nikel *et al*., 2014; Volke *et al*., 2021, 2022). These characteristics make *P. putida* a promising host for industrial bioconversion (Nikel & de Lorenzo, 2018; Poblete-Castro *et al*., 2012; Weimer *et al*., 2020).

As a proof-of-concept, we focussed on protocatechuate, the first common intermediate of *p*-coumarate, ferulate, vanillate, and vanillin degradation in *P. putida*. It is an important platform chemical for the production of a variety of aromatic compounds conventionally derived from petrochemical resources (Johnson *et al*., 2019; Kohlstedt *et al*., 2018; Notonier *et al*., 2021; Werner *et al*., 2023).

This study reports the co-substrate independent growth and protocatechuate production that solely rely on acetyl-CoA from the enoyl-CoA hydratase reaction in the *p*-coumarate and the ferulate branches of the lignin degradation pathway in *P. putida.* Furthermore, we investigate *in silico* the potential of heterologous C1 assimilation pathways for valorisation of the G lignin monomer, ferulate.

## Materials and Methods

### Bacterial Strains and Growth Media

Experiments were performed with *P. putida* EM42, a genome-reduced derivative of *P. putida* KT2440 (Martínez-García *et al*., 2014). Deletion of *pcaHG* in the EM42 strain was performed according to Volke *et al*. (2021) via homologous recombination of the I-SceI recognition site-containing suicide plasmid pSNW2-Δ*pcaHG* and subsequent curing with the I-SceI endonuclease expressing plasmid pQURE·6. Successful deletion mutants were identified via colony PCR. The plasmid for deletion of *pcaHG* was cloned in *Escherichia coli* DH5α λpir using USER cloning (Bitinaite *et al*., 2007). Generated strains, plasmids, and primer sequences are listed in the supplemental material (Supplementary Table S1).

For microtiter growth experiments and protocatechuate production experiments, two precultures were run prior to the main culture. The first preculture was grown overnight in LB medium, followed by a second overnight preculture in de Bont (dB) minimal medium (Hartmans *et al*., 1989) supplemented with 50 mM sodium acetate, 50 mM *p*-coumarate, or 50 mM ferulate, respectively. For main cultures, dB minimal medium supplemented with sodium acetate, *p*-coumarate, or ferulate was used at concentrations specified in the results section. The pH of *p*-coumarate and ferulate stocks was adjusted to 7.0 by addition of sodium hydroxide and the solutions were sterilised via filtration. Cultures with dB minimal medium were always supplemented with 10 g L^-1^ MOPS buffer (pH 7.0). All shake flask experiments were performed at 30 °C and 250 rpm in a rotary shaker.

### Microtiter Growth Experiments

Initial growth characterisation experiments were conducted using the Growth Profiler GP960 (EnzyScreen, Heemstede, The Netherlands). Per condition, 1 mL dB minimal medium was inoculated with an overnight preculture to an initial optical density at 600 nm (OD_600_) of 0.01. The inoculated culture was split into three single wells (300 μL each) in 96-well microtiter plates. The cultures were shaken at 250 rpm and at 30 °C. The cycle time was set to 600 s, and the shutter time was 4 ms. Green pixel values (G_value_) were obtained from photographed wells and converted to OD_600_ values via a polynomial regression (equation 1).

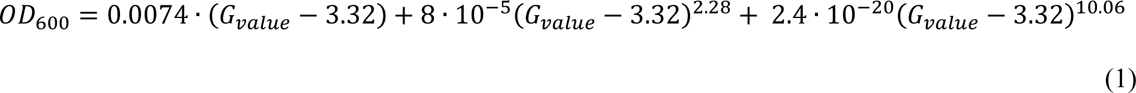

The equation was parameterised using *P. putida* EM42 cultures grown in LB medium with OD_600_ values ranging from 0.01 to 6.0 as determined by photometric analyses.

### Microbial Protocatechuate Production from *p*-Coumarate or Ferulate

Production of protocatechuate and other intermediate products downstream of the enoyl-CoA hydratase reaction was evaluated in shake flask experiments. For this, 25 mL of dB minimal medium with 75 mM *p*-coumarate / ferulate were inoculated to an initial OD_600_ of 0.01. Cultures were incubated at 250 rpm and 30 °C. For HPLC analysis, 0.5 mL cell suspension was withdrawn from the cultures throughout cultivation (see Figure 2) and centrifuged at 13,300 rpm for 1 min. 400 μL of the supernatant was transferred in a new tube and stored at -80 °C until further HPLC analysis.

### Quantification of Extracellular Aromatic Compounds

Supernatant samples were thawed and centrifuged at 16,000 g at 4 °C for 5 min to remove any remaining cells and cell debris. The supernatants were diluted appropriately prior to analysis. Samples were transferred to HPLC glass vials with conical glass inserts. The UHPLC analysis was performed on a ThermoFisher Scientific Vanquish DUO UHPLC with a DAD detector (ThermoFisher Scientific, Parkville, VIC, Australia). The compounds were separated on a Waters Acquity HSS T3 column (100×2.1 mm, 1.8 µm particles) (Waters, North Ryde, NSW, Australia). The column oven was set to 40 °C. A gradient was run for 11 min from 95 % 1:1000 formic acid in MilliQ water (mobile phase A) to 100 % 1:1000 formic acid in acetonitrile (mobile phase B) followed by 3 min flushing with 100 % mobile phase B. Between samples the column was equilibrated with 1:20 mobile phase B in mobile phase A for 6 min. The flow rate was set to 0.4 mL · min^-1^. Aromatic compounds were detected at 280 nm, 320 nm, and 360 nm.

Quantification was based on a standard calibration curve derived from serial dilution of chemical standards. The limits of quantification were 0.7 μM – 3 mM. Acquired data were analysed with the Chromeleon software 7.4 (ThermoFisher Scientific).

### Model Curation and Metabolic Modelling

A curated version of the most recent metabolic model of *P. putida* KT2440, *i*JN1463, (Nogales *et al*., 2020) was used. Reactions identified to enable growth on C1 compounds (CO_2_, formate) were set irreversible to prevent this growth phenotype nonnative to *P. putida* KT2440; these included reactions mediated by α-ketoglutarate dehydrogenase and phosphoribosylglycinamide formyltransferase (model reaction identifiers AKGDa and GARFT, respectively). Genes and corresponding reactions with pWW0 gene identifiers, originating from the TOL plasmid of *P. putida* mt-2, the progenitor strain of *P. putida* KT2440 (Bagdasarian *et al*., 1981) were removed, as well as Ni^2+^ requirement for biomass synthesis. A list of all curation steps can be found in Table 1.

**Table 1:**
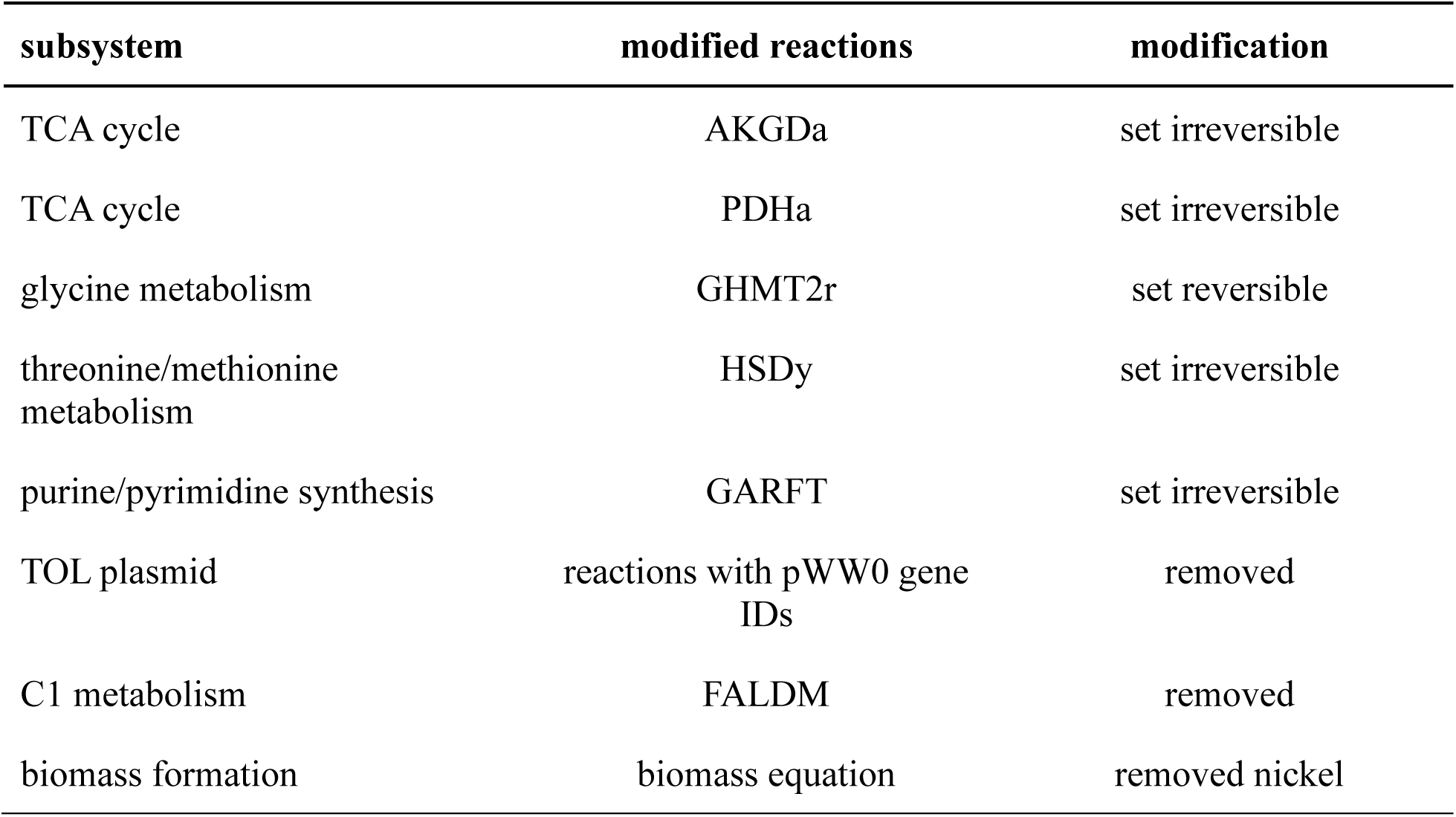
List of modifications for the curation of the genome-scale model *i*JN1463.

| subsystem | modified reactions | modification |
| --- | --- | --- |
| TCA cycle | AKGDa | set irreversible |
| TCA cycle | PDHa | set irreversible |
| glycine metabolism | GHMT2r | set reversible |
| threonine/methionine metabolism | HSDy | set irreversible |
| purine/pyrimidine synthesis | GARFT | set irreversible |
| TOL plasmid | reactions with pWW0 gene IDs | removed |
| C1 metabolism | FALDM | removed |
| biomass formation | biomass equation | removed nickel |

Flux balance analysis (FBA) was performed with COBRApy (Ebrahim *et al*., 2013). Uptake rates for simulations of growth on *p*-coumarate or ferulate were set as indicated in the respective section. Gene deletions were simulated using the COBRApy function ‘model.genes.id.knock_out()’, which sets the flux boundaries of all reactions associated with the respective gene to 0 mmol · gCDW ^-1^ · h^-1^. The objective function for all simulations was set to maximise biomass formation.

## Results and Discussion

### *In silico* Evaluation of Growth on C2 By-Product Fraction

Prior to *in vivo* experiments, growth on *p*-coumarate and ferulate was tested *in silico* using flux balance analysis (FBA). Simulations were run in COBRApy using a curated version of the published genome-scale model *i*JN1463 (Nogales *et al*., 2020). *p-*Coumarate and ferulate uptake rates were set to 4.5 mmol · g ^-1^ · h^-1^ for FBA, based on reported uptake rates of a range of aromatic compounds (Ravi *et al*., 2017). Protocatechuate production was simulated by deactivating the protocatechuate dioxygenase reaction PCADYOX, which oxidises protocatechuate to 3-carboxy-*cis*-muconate (Figure 1A). This reaction is associated with the genes *PP_4655* (*pcaH*) and *PP_4656* (*pcaG*), hence its inactivation resembles a Δ*pcaHG* deletion in *P. putida*. The uptake rates for all other carbon sources were set to 0 mmol g_CDW_^-1^ · h^-1^ forcing the model to rely solely on the available acetyl-CoA resulting from the enoyl-CoA hydratase reactions (COCOAHA and FCOAHA).

**Figure 1:**
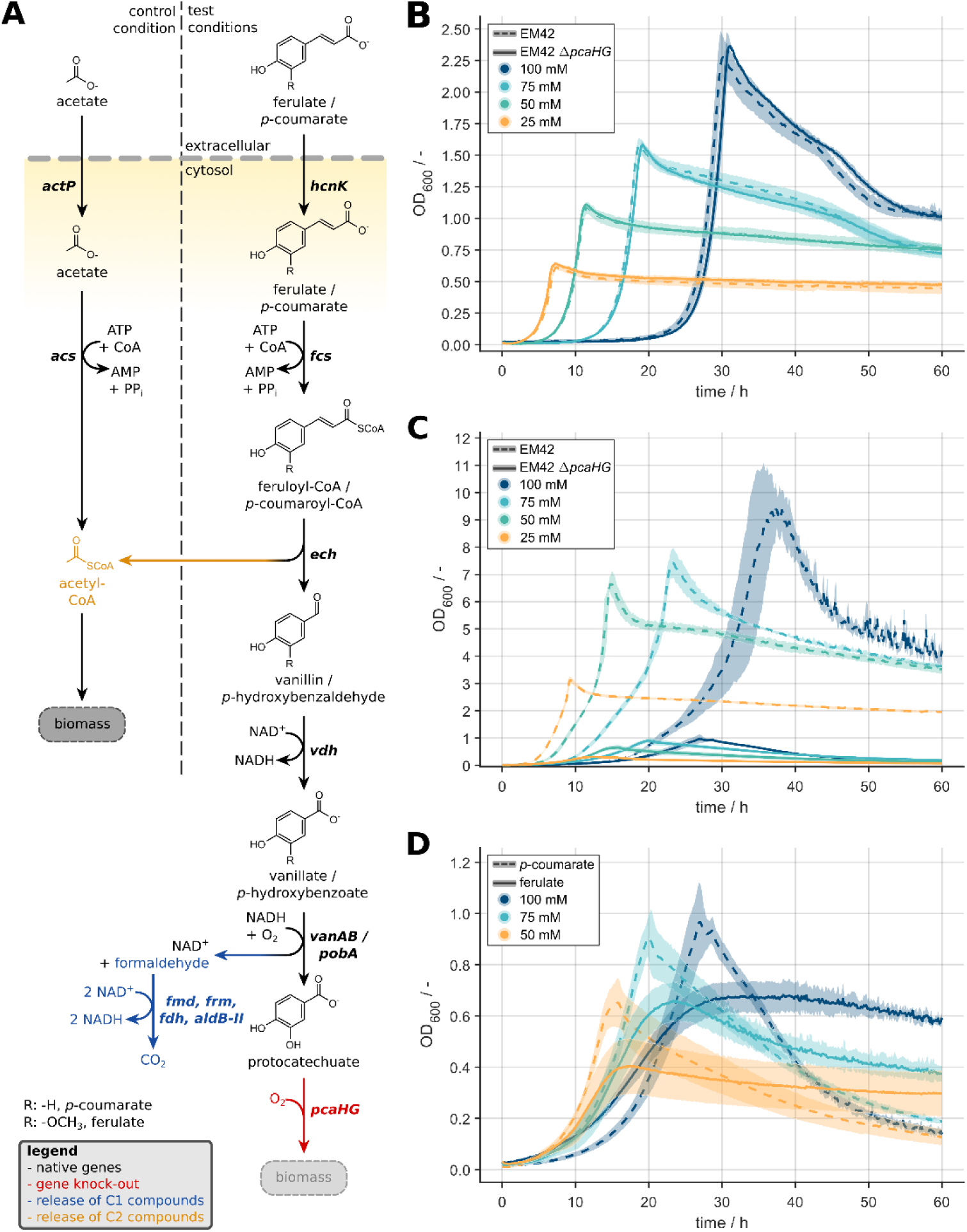
**A** Pathway for microbial conversion of *p*-coumarate/ferulate to protocatechuic acid in the Δ*pcaHG* mutant. The release of acetyl-CoA (from *p*-coumarate and ferulate) and formaldehyde (only from ferulate) is highlighted in orange and in blue, respectively. Subsequently, cleaved acetyl-CoA is the sole carbon source for biomass formation. Acetate conversion to acetyl-CoA is included for comparison. **B** Growth of EM42 (dashed line) and the Δ*pcaHG* deletion mutant (solid line) grown on different acetate concentrations. **C** Comparison of EM42 and the Δ*pcaHG* deletion mutant grown on different *p-*coumarate concentrations. **D** Comparison of the growth of the Δ*pcaHG* mutant on the by-product fractions from the degradation of *p*-coumarate (dashed lines) and ferulate (solid lines). The experimental error is included as the standard deviation of triplicate growth experiments and indicated by the shaded area.

The predicted biomass formation rates (Table 2) indicated that the amount of acetyl-CoA generated during conversion of *p*-coumarate or ferulate to protocatechuate is indeed sufficient for maintaining growth of a *pcaHG* deletion mutant incapable of further degrading protocatechuate as suggested by Salvachúa *et al*., (2018). As expected, the growth rates were highly decreased compared to the template model (-69 % for both substrates). Growth on ferulate led to a higher predicted growth rate than on *p-*coumarate (∼13 % for both cases, with and without deactivation of the protocatechuate dioxygenase). In contrast to *p-*coumarate, conversion of ferulate to protocatechuate produces formaldehyde in addition to acetyl-CoA (Figure 1A). While formaldehyde is not a carbon source of *P. putida*, full oxidation of this C1 compound via its formaldehyde and formate dehydrogenases provides additional NADH (2 molecules of NADH per molecule of formaldehyde; Roca *et al*., 2009; Roca & Ramos, 2009; Turlin & Puiggené *et al*., 2023; Zobel *et al*., 2017). Though it has not been reported that additional NADH leads to improved growth rates in *P. putida*, the biomass yield on the main carbon source can be increased by providing auxiliary co-substrates like formate serving as an electron source (Zobel *et al*., 2017). This benefit is based on reduced rates of substrate oxidation in the central carbon metabolism due to metabolic redirection from catabolism towards anabolism, which preserves the carbon from the main carbon source in biomass (Zobel *et al*., 2017). In the simulations, NADH produced from formaldehyde oxidation reduced the oxidation of acetyl-CoA in catabolic reactions, thereby increasing the carbon available for anabolic processes and the biomass formation rate.

**Table 2:** Predicted maximal growth rates on ferulate and *p*-coumarate of the template model and the model simulating deletion of *pcaHG* with and without enforced protocatechuate (PCA) production.

| biomass formation rate / h<sup>-1</sup> |  |  |  |
| --- | --- | --- | --- |
| substrate | template | $\Delta pcaHG$ | $\Delta pcaHG$ (enforced PCA production) |
| *p*-coumarate | 0.512 | 0.158 | 0.123 |
| ferulate | 0.581 | 0.178 | 0.178 |

The simulation with *p*-coumarate as the substrate did not show protocatechuate production. Instead, conversion stopped at *p*-hydroxybenzoate, the immediate precursor of protocatechuate. This occurred due to the NADH demand for the conversion of *p*-hydroxybenzoate to protocatechuate. NADH formed in the previous step (the conversion of *p*-hydroxybenzaldehyde to *p-*hydroxybenzoate) could be reinvested here. However, since the simulations maximised growth, NADH is preferentially used for biomass formation, which is limiting in this scenario where only the acetyl-CoA by-product fraction of the *p*-coumarate branch is available. Enforcing protocatechuate production in the simulation with *p*-coumarate, lowered the biomass formation rate by 22 % (Table 2). Similar to the formation of protocatechuate from *p*-hydroxybenzoate, the production of protocatechuate from vanillate in the ferulate branch requires one molecule of NADH per molecule of vanillate. However, the oxidation of the released formaldehyde provides additional reduced redox equivalents that outweigh the NADH demand required for vanillate conversion. Nevertheless, no complete conversion of vanillate to protocatechuate was observed in the simulations with protocatechuate formation rates reaching only 73 % of the ferulate uptake rate. However, this is a modelling artefact and not due to a lack of energy, as enforcement of protocatechuate production from ferulate did not lead to a reduction in biomass formation suggesting that conversion of vanillate to protocatechuate potentially leads to excess NADH.

Indeed, the limitation by the last step in protocatechuate production from *p*-coumarate/ferulate as suggested by the model has previously been observed *in vivo* (Kuatsjah *et al*., 2022; Mohamed *et al*., 2020; Werner *et al*., 2023).

### *In vivo* Growth on C2 By-Product Fraction from *p*-Coumarate/Ferulate Degradation

To confirm the *in silico* results, microtiter growth experiments with *p*-coumarate and ferulate were performed. Acetate was included as the control condition. The activation of acetate to acetyl-CoA requires the same co-factors as the activation of *p*-coumarate and ferulate (Figure 1A). The remaining two steps for protocatechuate production catalysed by vanillin dehydrogenase and *p-*hydroxybenzoate dehydrogenase, encoded by *vnd* and *pob*A respectively, are redox balanced. Therefore, they do not increase the net cofactor demand. Hence, growth on acetate should be similar to *p*-coumarate in a strain incapable of performing protocatechuate catabolism.

Initial comparison of the growth behaviour of the parental strain *P. putida* EM42 (EM42) and the Δ*pcaHG* mutant on acetate revealed identical growth of both strains (Figure 1B and Table 3) showing that the deletion of *pcaHG* does not affect acetate metabolism. On *p*-coumarate, EM42 Δ*pcaHG* reached a maximum optical density (OD_max_) of only 9.6–12.4 % of the maximum OD_600_ of the parental strain (Figure 1C and Table 3). This was expected due to the smaller proportion of carbon available for growth. Unlike EM42, the *pcaHG* deletion mutant can only utilise the C2 by-product fraction (acetyl-CoA) lowering the available carbon pool by ∼80 %.

**Table 3:** Comparison of maximum growth rate (μ_max_) and maximum optical density (OD_max_) of EM42 and EM42 Δ*pcaHG* grown on different concentrations of acetate and *p*-coumarate. Errors are based on the standard deviation of biological triplicates.

| strain | substrate | concentration / mM | $\mu_{\max}$ / h<sup>-1</sup> | $OD_{\max}$ / - |
| --- | --- | --- | --- | --- |
| EM42 | acetate | 25 | 0.717±0.029 | 0.61±0.02 |
|  |  | 50 | 0.622±0.043 | 1.08±0.04 |
|  |  | 75 | 0.536±0.007 | 1.58±0.05 |
|  |  | 100 | 0.478±0.015 | 2.29±0.16 |
|  | *p*-coumarate | 25 | 0.871±0.074 | 3.12±0.11 |
|  |  | 50 | 0.691±0.024 | 6.77±0.25 |
|  |  | 75 | 0.446±0.030 | 7.43±0.53 |
|  |  | 100 | 0.301±0.027 | 10.24±0.88 |
|  | ferulate | 25 | 0.782±0.097 | 2.66±0.26 |
|  |  | 50 | 0.592±0.019 | 5.41±0.10 |
|  |  | 75 | 0.464±0.017 | 7.69±0.34 |
|  |  | 100 | 0.285±0.074 | 5.22±0.04 |
| EM42 $\Delta pcaHG$ | acetate | 25 | 0.760±0.014 | 0.64±0.01 |
|  |  | 50 | 0.643±0.018 | 1.11±0.01 |
|  |  | 75 | 0.508±0.007 | 1.59±0.03 |
|  |  | 100 | 0.472±0.005 | 2.4±0.06 |
|  | *p*-coumarate | 25 | 0.370±0.020 | 0.30±0.01 |
|  |  | 50 | 0.380±0.030 | 0.65±0.09 |
|  |  | 75 | 0.267±0.042 | 0.92±0.10 |
|  |  | 100 | 0.192±0.020 | 1.03±0.07 |
|  | ferulate | 25 | 0.290±0.046 | 0.15±0.05 |
|  |  | 50 | 0.252±0.038 | 0.41±0.11 |
|  |  | 75 | 0.251±0.017 | 0.66±0.08 |
|  |  | 100 | 0.181±0.005 | 0.70±0.05 |

Despite the same cofactor demand, growth on acetyl-CoA derived from *p-*coumarate led to 60 % lower final optical densities compared to growth on acetate, and growth was also slowed by 40–60 %. These differences could be attributed to the toxicity of *p*-coumarate and its aromatic breakdown products (Calero *et al*., 2018; Mohamed *et al*., 2020; Salvachúa *et al*., 2018; Zhang *et al*., 2021).

Next, we compared growth of the *pcaHG* deletion mutant on *p*-coumarate and ferulate, (Figure 1D), provided at concentrations between 25 and 100 mM. While the lower substrate concentrations resulted in the highest observed growth rates on the byproduct fractions of both substrates, with the cultures on 25 mM and 50 mM *p*-coumarate exceeding the cultures on 25 mM and 50 mM ferulate, at 100 mM no significant difference was observed (Table 3). Growth on *p*-coumarate resulted in higher maximal optical densities under all tested conditions. It was anticipated that formaldehyde released from ferulate should lead to higher final biomass concentrations similar to the results of co-feeding glucose and formate to *P. putida* KT2440 (Zobel *et al*., 2017).

### Quantification of Substrate Conversion to Protocatechuate

The observed growth behaviour of the deletion mutant on ferulate indicated a limitation in either formaldehyde oxidation as reported for *P. putida* KT2440 grown on vanillate or in the demethylation of vanillate and hence formaldehyde release. Indeed, it was shown previously that *P. putida*’s native vanillate O-demethylase (encoded by *vanAB*) can constitute a bottleneck in ferulate conversion (Salvachúa *et al*., 2018; Werner *et al*., 2023). In both cases, formaldehyde cannot be used as an electron source.

To test these hypotheses, by-product formation was determined in shake flask experiments with 75 mM *p*-coumarate or ferulate (Figure 2). Under both conditions, the aromatic acid precursors of protocatechuate, *p*-hydroxybenzoate and vanillate, accumulated, as reported by Werner *et al*. (2023). Accumulation of vanillate in the ferulate conditions limited the release of formaldehyde and NADH generation from its oxidation, explaining the absence of a growth advantage over *p*-coumarate. Furthermore, we observed the accumulation of *p*-hydroxybenzoate in the *p*-coumarate condition, which could be attributed to the beneficial NADH savings suggested by the *in silico* results. No accumulation of the aldehyde products (*p*-hydroxybenzaldehyde/vanillin) was observed.

**Figure 2:**
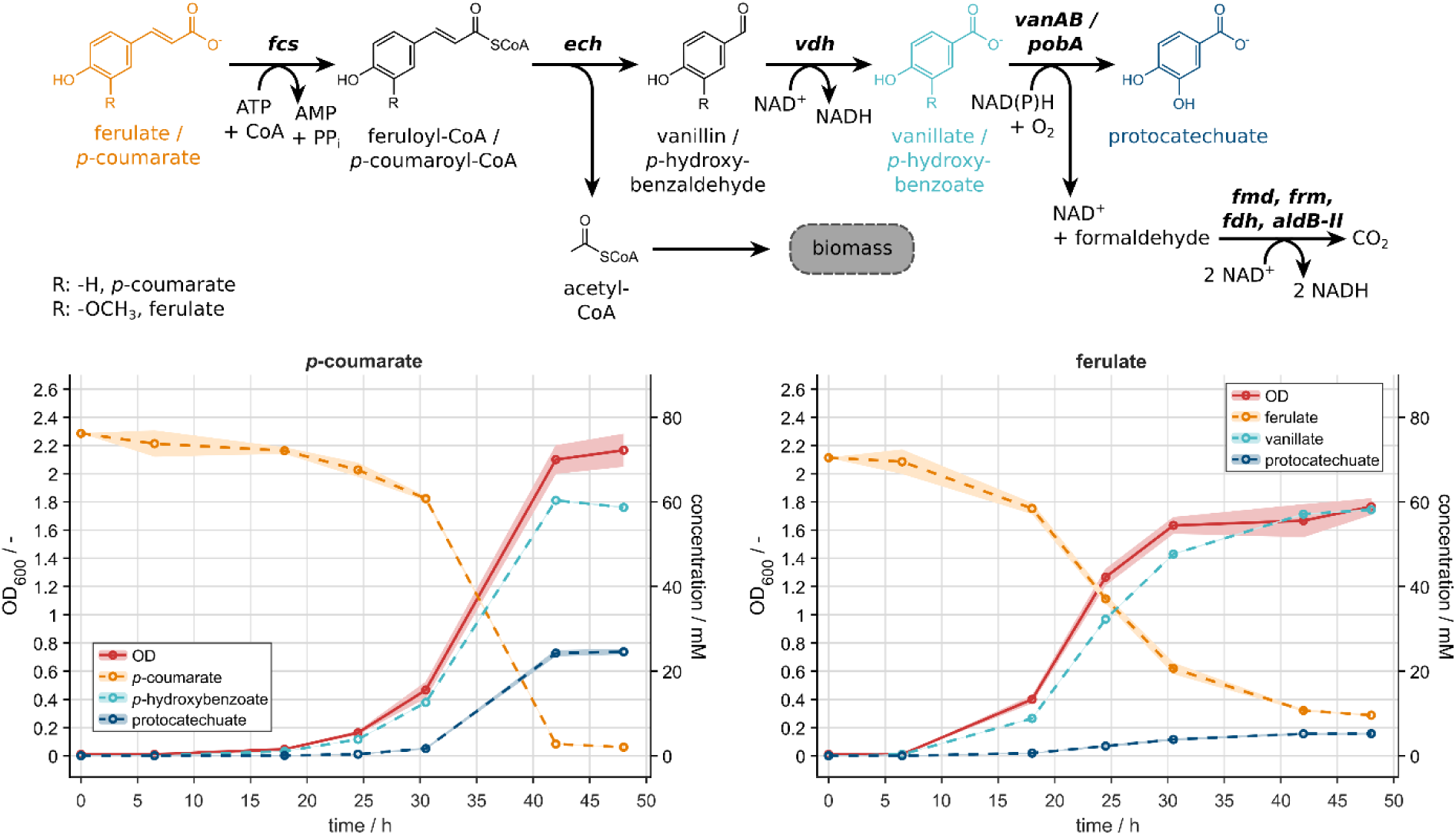
**A** Pathway for protocatechuate production from *p*-coumarate/ferulate in the Δ*pcaHG* mutant. The detected compounds ferulate/*p*-coumarate, vanillate/*p*-hydroxybenzoate, and protocatechuate are highlighted in orange, light blue, and dark blue, respectively. **B** Concentration profile of *p*-coumarate, *p*-hydroxybenzoate, and protocatechuate of the Δ*pcaHG* mutant grown on *p*-coumarate. **C** Concentration profile of ferulate, vanillin, and protocatechuate of the Δ*pcaHG* mutant grown on ferulate. Aromatic compound concentrations are shown as dashed lines, and the measured OD_600_ is included as solid line. The standard deviation of triplicates is indicated by the shaded area.

Overexpressing the native *pobA* and *vanAB* genes or their homologs, can support *p-*hydroxybenzoate and vanillate conversion (Kuatsjah *et al*., 2022; Werner *et al*., 2023), with the latter leading to formaldehyde release. *P. putida* possesses a set of formaldehyde and formate dehydrogenases with high activities (Roca *et al*., 2009; Turlin *et al*., 2023). Despite its arsenal of formaldehyde and formate dehydrogenases constitutively or conditionally expressed to prevent toxicity (Roca & Ramos, 2009; Turlin et al., 2023), formaldehyde accumulation was observed during growth on vanillate (Lee et al., 2021, 2022). To observe the beneficial effects of formaldehyde oxidation predicted by metabolic modelling, stronger constitutive expression of formaldehyde and formate dehydrogenases might be necessary.

Product yields were compared under both conditions to assess overall conversion efficiency. Only 32.3±0.8 % of *p*-coumarate and 7.4±0.2 % of ferulate were converted to protocatechuate. The rest accumulated as *p*-hydroxybenzoate (77.0±0.8 %) and vanillate (82.6±2.0 %), while 2.7±0.0 % of *p*-coumarate and 13.6±0.1 % ferulate remained in the media at the end of the cultivation. The sum of the final concentrations of protocatechuate and the respective aromatic acid precursor indicated a molar yield of approx. 100 % in the case of *p*-coumarate and 90.0±2.2 % for ferulate.

Previous studies reported molar yields of 93±1 % and 94±7 % for ketoadipate production from *p*-coumarate/ferulate (Werner *et al*., 2023) but supplemented additional carbon sources like glucose or acetate for biomass formation (Becker *et al*., 2018; Johnson *et al*., 2017; Werner *et al*., 2023). When accounting for this additional carbon, the overall carbon yield was only 52.6 % in the case of the study of Werner *et al*. (2023). In comparison, we achieved a carbon yield of 78 % for the sum of products, confirming that the C2 by-product fraction alone is sufficient for maintaining the growth of lignin valorisation strains while increasing the carbon efficiency. Indeed, the total product yield equals the theoretical maximum carbon yield for *p*-coumarate.

### *In silico* Study of C1 Assimilation Pathways for Application in Lignin Valorisation

To lower CO_2_ emissions from formaldehyde oxidation and increase the carbon available for growth, integration of C1 assimilation pathways into lignin valorisation strains can be considered. Indeed, overexpression of key enzymes of the ribulose-5-monophosphate pathway in lignin-degrading bacteria with native 3-hexulose-6-phosphate-synthase and the 6-phosphate-3-hexulose-isomerase, has improved growth on vanillate (Mitsui *et al*., 2003). Including C1 assimilation pathways may also aid the growth of the Δ*pcaHG* mutant. Recent advances in heterologous expression of C1 assimilation pathways in heterotrophic microorganisms *demonstrate* its feasibility (Bruinsma *et al*., 2023; He *et al*., 2020, 2020; Keller *et al*., 2022; Lu *et al*., 2019; Turlin *et al*., 2022; Wenk *et al*., 2022).

To study the effect of C1 assimilation pathways on lignin valorisation with *P. putida*, we compared different C1 assimilation pathways *in silico*, the synthetic acetyl-CoA (SACA) pathway, ribulose-5-monophosphate (RuMP) cycle, reductive glycine (RG) pathway, and the homoserine (HOM) cycle (see Figure 3). These pathways were chosen based on their different architecture, different co-factor demands, and the different entry points of their output metabolites into the central carbon metabolism. Simulations were carried out with the curated genome-scale model *i*JN1463 in COBRApy extended with the reactions for the C1 assimilation pathways. The sole carbon source for the simulations was ferulate, set at an uptake rate of 4.5 mmol · g_CDW_ ^-1^ · h^-1^. Simulations with *p*-coumarate were included for comparison.

**Figure 3:**
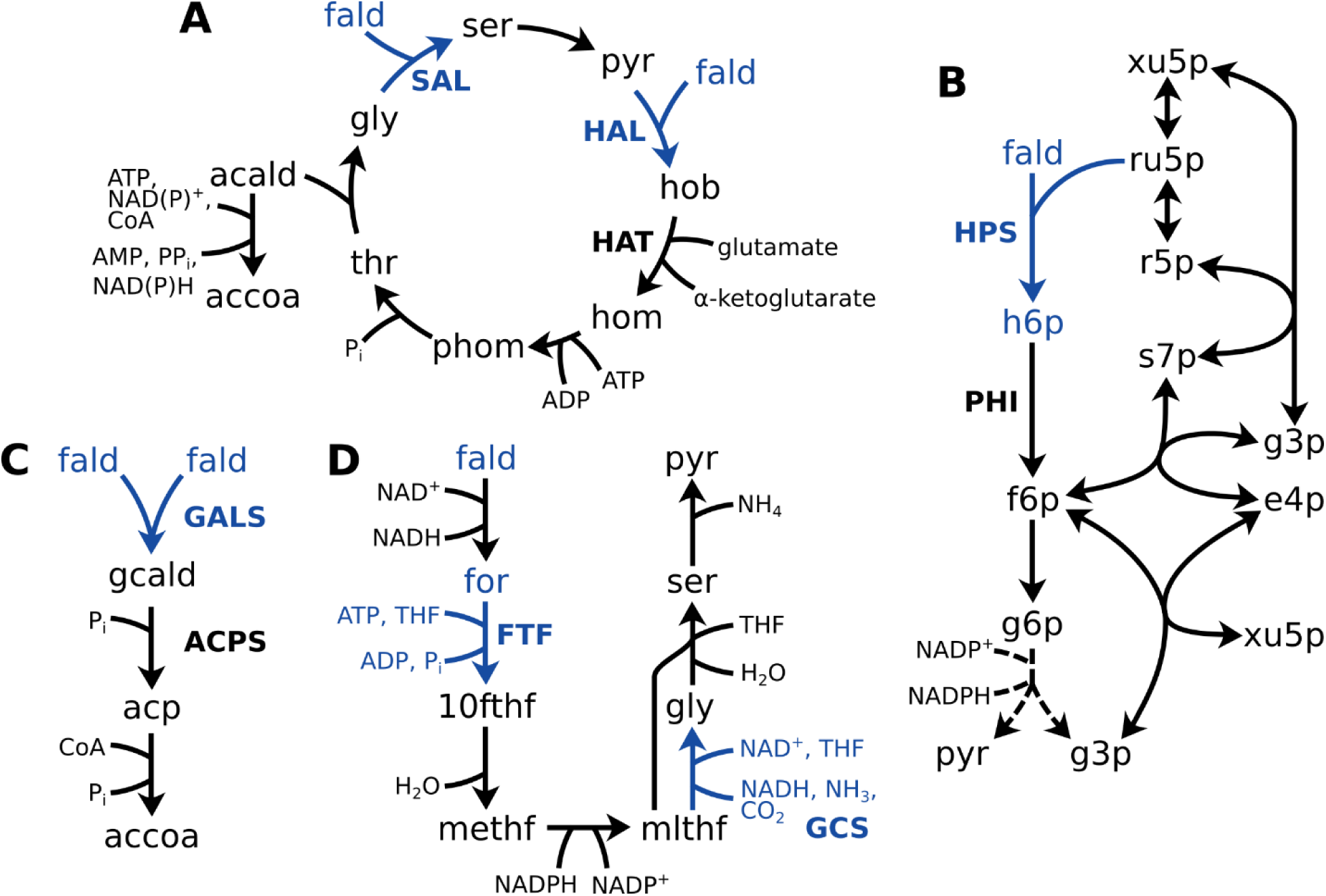
C1 assimilation pathways for the *in silico* study on the application of C1 assimilation in lignin valorisation system: **A** homoserine (HOM) cycle (He *et al*., 2020); **B** ribulose-5-monophosphate (RuMP) cycle (Bar-Even *et al*., 2013); **C** synthetic acetyl-CoA (SACA) pathway (Lu *et al*., 2019); **D** reductive glycine (RG) pathway (Bar-Even *et al*., 2013; Bruinsma *et al*., 2023; Turlin *et al*., 2022). Metabolites and co-factors – fald: formaldehyde, gcald: glycolaldehyde, acp: acetyl-phosphate, accoa: acetyl-CoA, ru5p: ribulose-5-monophosphate, h6p: hexulose-6-phosphate, f6p: fructose-6-phosphate, pyr: pyruvate, g6p: glucose-6-phosphate, 6pgl: 6-phosphoglucono-1,5-lactone, 6pgc: 6-phosphogluconate, for: formate, 10fthf: 10-formyl-THF, methf: 5,10-methenyl-THF, mlthf: 5,10-methylene-THF, gly: glycine, ser: serine, hob: 4-hydroxy-2-oxobutanoate, hom: homoserine, phom: phosphohomoserine, thr: threonine, acald: acetaldehyde, THF: tetrahydrofolate. Enzymes – GALS: glycolaldehyde synthetase, ACPS: acetylphosphate synthetase, H6PS: hexulose-6-phosphate synthetase, H6PI: hexulose-6-phosphate isomerase, FTF: formate-THF ligase, GCS: glycine cleavage system, SAL: serine aldolase, HAL: 4-hydroxy-2-oxobutanoate aldolase, HAT: 4-hydroxy-2-oxobutanoate aminotransferase,

All simulations suggested that assimilation of formaldehyde is beneficial for the growth of the lignin valorisation strain, with the RuMP cycle enabling the highest improvement of the biomass formation rate of 11 % (Table 4). While the SACA pathway and the RuMP pathway assimilated all formaldehyde, the CO_2_ emission rates indicated that the assimilated carbon was oxidised in the central carbon metabolism. Overall, the benefit of C1 assimilation was low due to the energy demand of the C1 assimilation.

**Table 4:** Simulation of growth of a protocatechuate production model (Δ*pcaHG*) on the C2 and C1 by-product fractions from ferulate using different C1 assimilation pathways. The results for the Δ*pcaHG* model with *p*-coumarate (*p*CA) and ferulate are included for comparison. Biomass formation rate, percentage of not assimilated, fully oxidised C1 fraction (C1_ox_), and CO_2_ emission rate are listed. SACA: synthetic acetyl-CoA pathway; RuMP: ribulose-5-monophosphate cycle; RG: reductive glycine pathway; HOM: homoserine cycle.

| model | $\mu / \text{h}^{-1}$ | C1<sub>ox</sub> / % | CO<sub>2</sub> emission / $\text{mmol} \cdot \text{g}_{\text{CDW}}^{-1} \cdot \text{h}^{-1}$ |
| --- | --- | --- | --- |
| $\Delta pcaHG$ (*pCA*) | 0.158 | — | 2.65 |
| $\Delta pcaHG$ | 0.178 | 98.4 | 6.34 |
| SACA | 0.194 | 0.0 | 5.67 |
| RuMP | 0.198 | 0.0 | 5.54 |
| RG | 0.192 | 87.3 | 5.77 |
| HOM | 0.191 | 45.7 | 5.82 |

C1 assimilation pathways are highly energy demanding. Thus, photo-/chemoautotroph microbes that natively use these pathways rely on additional energy sources, such as light, hydrogen, phosphite, and auxiliary C1 co-substrates or reduced C1 sources like carbon monoxide (Claassens *et al*., 2018; Figueroa *et al*., 2018). While sacrificial C1 compounds have been shown to be promising electron donors for heterotrophic microbes (Turlin *et al*., 2022; Zobel *et al*., 2017), the aim of this study was to increase the carbon yield of the proposed lignin valorisation strain design. Therefore, we decided to analyse the effect of including a carbon-free electron carrier in the models that enables electron supply from, e.g., hydrogen or phosphite on fuelling formaldehyde assimilation. Hydrogen is the global focus for several different applications, and research and innovation for sustainable production of green hydrogen as an energy carrier is vigorously pursued.

Heterologous expression of an oxygen-tolerant hydrogenase from *Cupriavidus necator* enabled hydrogen oxidation in *P. putida* and the augmented redox power was shown to improve the NADH-dependent biotransformation of *n*-octane to 1-octanol (Lonsdale *et al*., 2015). Similar to hydrogen, phosphite can be produced sustainably (Zhai *et al*., 2022), and it can serve as phosphate source in *P. putida* when expressing a phosphite dehydrogenase, providing one molecule of NADH per molecule of phosphite (Asin-Garcia *et al*., 2022). Furthermore, phosphite has a very low reduction level (-650 mV), making the phosphite dehydrogenase an attractive candidate for heterologous carbon-free NADH regeneration (Claassens *et al*., 2018; Costas *et al*., 2001).

The results of the simulations were identical for hydrogen and phosphite as electron donors and are detailed for hydrogen supply in Table 5. However, in the case of phosphite, substantial phosphate export was observed due to a up to 100-fold higher demand for NADH than for phosphate.

**Table 5:** Simulation of growth on the C2 and C1 by-product fractions from ferulate of a protocatechuate production model (Δ*pcaHG*) complemented with different C1 assimilation pathways and a hydrogenase reaction. The results for the Δ*pcaHG* model with *p*-coumarate (*p*CA) and ferulate are included for comparison. Biomass formation rate, percentage of not assimilated, fully oxidised C1 fraction (C1_ox_), and CO_2_ emission rate are listed. SACA: synthetic acetyl-CoA pathway; RuMP: ribulose-5-monophosphate cycle; RG: reductive glycine pathway; HOM: homoserine cycle.

| model | $\mu / h^{-1}$ | $C1_{ox} / \%$ | $CO_2$ emission / $mmol \cdot g_{CDW}^{-1} \cdot h^{-1}$ |
| --- | --- | --- | --- |
| $\Delta pcaHG$ (pCA) | 0.175 | – | 1.94 |
| $\Delta pcaHG$ | 0.178 | 93.8 | 6.13 |
| SACA | 0.265 | 0.0 | 2.84 |
| RuMP | 0.293 | 0.0 | 1.69 |
| RG | 0.350 | 0.0 | -0.60 |
| HOM | 0.350 | 0.0 | -0.60 |

The utilisation of the inorganic electron source abolished formaldehyde oxidation in all models augmented with C1 assimilation pathways. CO_2_ emission rates were reduced by 54 % and 72 % for the SACA pathway and RuMP cycle, respectively. In the case of the RG pathway and the HOM cycle, the simulations suggested CO_2_ utilisation leading to net CO_2_ uptake instead of CO_2_ production. For the model incorporating the RG pathway, CO_2_ assimilation was based on the reductive glycine complex, which fixes one molecule of CO_2_ per molecule of glycine formed. Within the HOM cycle, increased fluxes through the pyruvate carboxylase led to CO_2_ assimilation, whereby one molecule of CO_2_ is condensed onto pyruvate to generate one molecule of oxaloacetate.

These *in silico* results demonstrate potential benefits of co-assimilating lignin-derived formaldehyde with C2 by-product fractions from lignin aromatic monomers to maximally conserve the monomeric lignin substrates into the desired product.

### Model-Based Identification of a Novel C1 Assimilation Pathway for Growth-Coupled Lignin Valorisation

The simulations involving the HOM cycle relied on the activity of the pyruvate carboxylase, an enzyme that was shown to carry significant flux in *P. putida* (Zobel *et al*., 2017). In addition, the simulations revealed that key reactions of the HOM cycle, namely the 4-hydroxy-2-oxobutanoate aldolase and aminotransferase reactions were inactive. Instead, the model used an alternative C1 assimilation pathway. This novel pathway is a hybrid that utilises the serine aldolase (SAL) reaction from the HOM cycle to condense formaldehyde to glycine, generating serine, and integrates parts of the serine threonine (ST) cycle, a formate assimilation pathway described by Wenk *et al*. (2022) (see Figure 4A). In the ST cycle, formate is conjugated to tetrahydrofolate (THF) to form methylene-THF, which transfer the C1 moiety onto glycine, producing serine. Deamination of serine yields pyruvate, the substrate for pyruvate carboxylase, which fixes CO_2_-derived bicarbonate (HCO_3_^-^) resulting in oxaloacetate. Oxaloacetate is further converted to threonine, which when cleaved by threonine aldolase, regenerates glycine, thereby closing the C1 assimilation pathway, and releases the C2 product acetaldehyde. Except for the assimilation of formate, all reactions of the ST cycle are native to *P. putida*, while serine aldolase activity has been shown to be functional in *P. putida* with the threonine aldolase native to *Escherichia coli* (Puiggené *et al*., 2025). Introducing a single gene encoding a serine aldolase together with the NADH regeneration module (hydrogenase/phosphite operon) in *P. putida* would enable formaldehyde co-assimilation via this novel C1 assimilation pathway, subsequently referred to as SAL shunt.

**Figure 4:**
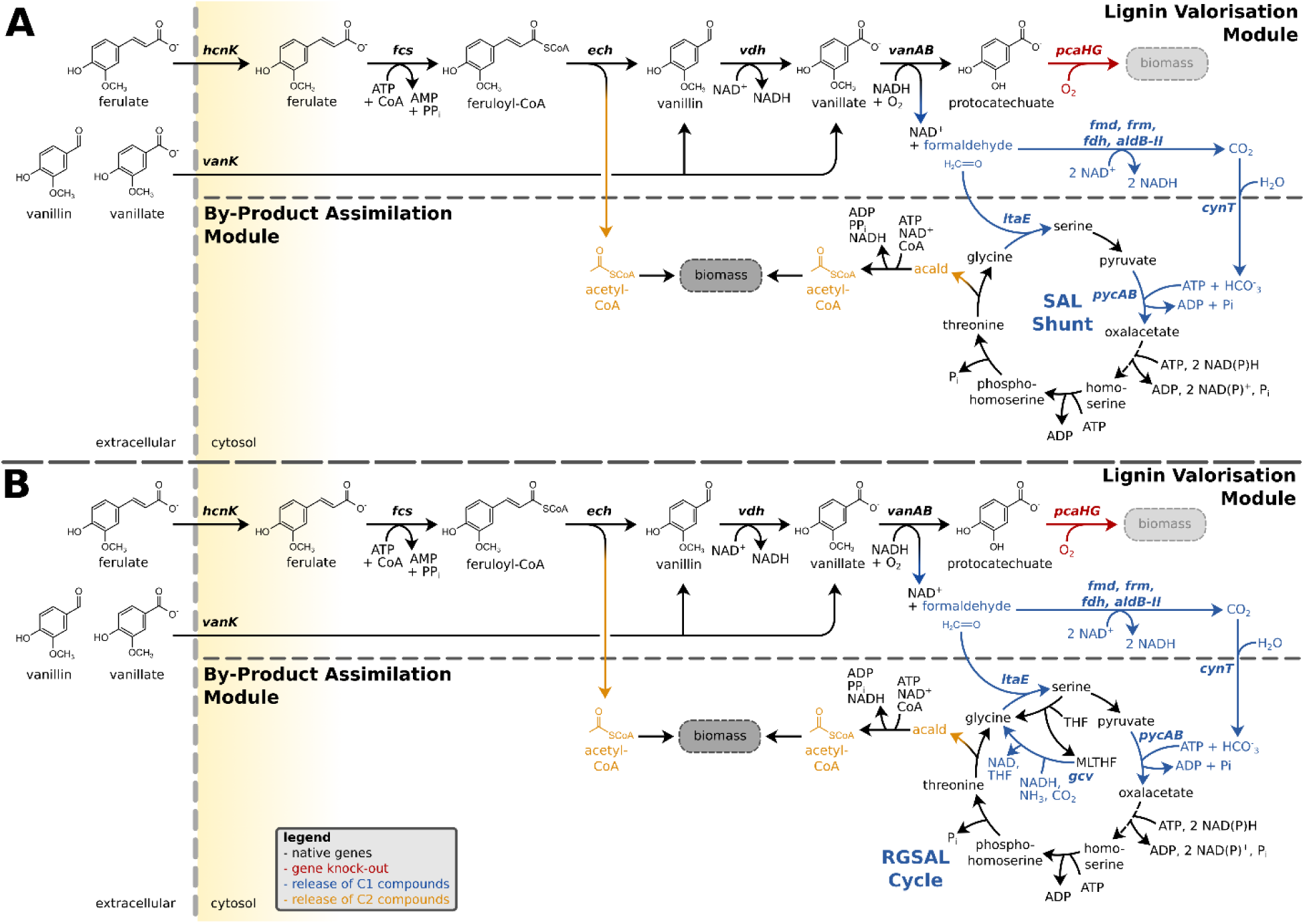
Novel C1 assimilation pathways proposed in a protocatechuate-producing strain, enabling capture of the C1 by-product fraction from G lignin monomers (vanillin, vanillate, ferulate); **A** via the serine aldolase (SAL) shunt; **B** via the reductive glycine pathway-serine aldolase hybrid cycle (RGSAL).

Considering the improvement in simulated biomass formation and the dependence on a single non-native reaction for formaldehyde assimilation while also fixing CO_2_, the SAL shunt was identified to be the most suitable C1 assimilation pathway among the tested pathways for the application in lignin valorisation strains solely relying on the C1 and C2 by-product fractions for growth.

In addition, we performed a competitive FBA with a model including reactions of all tested C1 assimilation pathways to identify other potential combinations of C1 assimilating reactions, similar to the SAL shunt. The model suggested a pathway combining the SAL shunt with the RG pathway (see Figure 4B). This pathway assimilates formaldehyde and CO_2_ in a circular fashion. First formaldehyde is assimilated via the serine aldolase generating serine. A smaller fraction of serine is deaminated to pyruvate, which then captures CO_2_ via the pyruvate carboxylase as in the SAL shunt; serine is cleaved via the reversible glycine hydroxymethyl transferase (encoded by *glyA*I-II) into glycine and 5,10-methylene-THF. The latter serves as a substrate for the reductive glycine complex of the RG pathway capturing one molecule of CO_2_ and producing glycine. The pathway has thus three streams for glycine generation: 1) via serine cleavage by glycine hydroxymethyl transferase activity, 2) via the reductive glycine complex, and 3) via threonine cleavage catalysed by the threonine aldolase .

While no benefit in terms of biomass formation was computed, this pathway has a significantly reduced NADH demand in comparison to the SAL shunt and the RG pathway. The required NADH per assimilated carbon atom is 1.64 for the SAL shunt and 0.52 for the RG pathway. The RGSAL cycle requires only 0.33 NADH per assimilated carbon atom. In all cases, growth of the lignin valorisation strain would be coupled to C1 and C2 by-product formation and therefore to the production of protocatechuate from methoxylated lignin monomers.

## Conclusion

This study demonstrates a novel approach to lignin valorisation, highlighting the feasibility of utilising C2 (acetyl-CoA) by-products from *p*-coumarate and ferulate degradation for growth while conserving the valuable aromatic core of lignin monomers in the target product. Using a *P. putida* EM42 *pcaHG* deletion mutant, a near-perfect molar yield of approximately 100% was achieved for ferulate and *p*-coumarate conversion with 78% of the substrate carbon conserved in the aromatic degradation products. This represents the highest reported carbon yield for biotechnological lignin utilization to date.

Our *in silico* analyses showed limited potential for capturing formaldehyde) from methoxylated lignin monomers as additional carbon source due to energy limitation. However, when enabling the utilisation of electron donors such as hydrogen or phosphite, simulations not only identified novel C1 assimilation pathway variants but also suggested a significant boost in carbon efficiency and reduced carbon loss as CO_2_.

In conclusion, our study suggests a promising glucose-free lignin valorisation strategy solely relying on C1 and C2 by-product co-assimilation for growth enabled by synthetic C1 assimilation pathways.

## Acknowledgments

The authors would like to thank the other members of the Systems Environmental Microbiology group at DTU Biosustain (Denmark) for their support and guidance in genetic engineering of *Pseudomonas putida* and their expertise in C1 assimilation pathways provided.

The authors thank Dr Gabi Netzel of the Queensland Node of Metabolomics and Proteomics Australia for their support and assistance in this work. Q-MAP is supported by Bioplatforms Australia, an NCRIS funded initiative.

