## Supplementary Table S1 for "Co-Substrate Free Valorisation of Lignin Monomers by Assimilation of C1 and C2 By-Products"

### Supplemental Materials

**Table S1:** List of strains, primers, and plasmids used in this study.

| **strains** | **description** | **references** |
|---|---|---|
| *E. coli* DH5α λpir | cloning host for construction of deletion plasmids carrying the R6K ori | Platt *et al*., 2000 |
| *E. coli* DH5α pQURE | host carrying the plasmid for curation of pEMG derivatives containing a I-SceI recognition site | Volke *et al*., 2021 |
| *E. coli* DH5α λpir pSNW2-Δ*pcaHG* | cloning host carrying the suicide plasmid for deletion of the genes *pcaH* and *pcaG* in *P. putida* EM42 via homologous recombination | this study |
| *P. putida* EM42 | template strain of *P. putida*, a genome-reduced variant of *P. putida* strain KT2440 | Martínez-García *et al*., 2014 |
| *P. putida* EM42 Δ*pcaHG* | *pcaHG* deletion mutant of *P. putida* EM42 | this study |

| **primers** | **sequences** | **description** | **references** |
|---|---|---|---|
| pcaHG_HR1_fw | AGATCCUCGTCGGTCAATGCCGCCC | fw-primer for amplification of the homologous 500 bp region upstream of the *pcaHG* locus | this study |
| pcaHG_HR1_rv | ATGTGAGGUGAAGCTTGGGGCCGCTCT | rv-primer for amplification of the homologous 500 bp region upstream of the *pcaHG* locus | this study |
| pcaHG_HR2_fw | ACCTCACAUGCCGGTTTCCTCTCTTGGAATTGT | fw-primer for amplification of the homologous 500 bp region downstream of the *pcaHG* locus | this study |
| pcaHG_HR2_rv | AGGTCGACUCGGCCTGGGCGAACAGGG | rv-primer for amplification of the homologous 500 bp region downstream of the *pcaHG* locus | this study |
| pSNW-USER_F | AGTCGACCUGCAGGCATGCAAGCTTCT | fw-primer for linearisation of the pSNW backbone | Volke *et al*., 2021 |
| pSNW-USER_R | AGGATCUAGAGGATCCCCGGGTACCG | rv-primer for linearisation of the pSNW backbone | Volke *et al*., 2021 |

| **plasmids** | **description** | **reference** |
|---|---|---|
| pSNW2 | derivative of pEMG, suicide plasmid for gene deletions via homologous recombination, $Km^R$ | Volke *et al*., 2021 |
| pQURE·6 | plasmid for curation genomically integrated suicide plasmids containing I-SceI recognition sites, $Gm^R$ | (Volke *et al*., 2021) |
| pSNW2-Δ*pcaHG* | suicide plasmid for deletion of the *pcaHG* locus in *P. putida* EM42, $Km^R$ | this study |
